# Histological divergence underlying globular body shapes in ornamental goldfish

**DOI:** 10.1101/2025.10.01.679695

**Authors:** Kinya G Ota, Gembu Abe, Chen-Yi Wang, Ing-Jia Li, Paul Gerald Layague Sanchez

## Abstract

Body shape diversity in vertebrates reflects a complex interplay between functional demands, environmental constraints, and internal developmental mechanisms. Various environments have promoted diverse morphological adaptations not only under natural but also domesticated conditions. One of the most remarkable examples of artificially induced morphology is found in the domesticated ornamental goldfish (*Carassius auratus*), which has diversified into numerous strains with strikingly different body shapes through prolonged human selection. In this study, we compared the body shapes of representative goldfish strains: the single-tail common goldfish (wild-type), *Ryukin, Oranda, Pearl scale*, and *Ranchu*. Our analysis revealed that the *Ryukin* and *Pearl scale* strains exhibit significantly greater body circularity in dorsal view compared to the other strains. Further anatomical and histological analyses showed that *Pearl scale* goldfish possess a thicker lateral body wall along with increased adipose tissue accumulation and reduced muscle fiber density, unlike *Ryukin* goldfish. These findings suggest that similar globular body shapes in different goldfish strains have arisen through distinct developmental pathways, exemplifying morphological convergence accompanied by histological divergence. We further discuss adipose accumulation in *Pearl scale* goldfish in relation to natural examples, providing insight into how function, morphology, and tissue organization may be interlinked in the evolution of globular body shapes.

## BACKGROUND

The diversity of body shapes observed in aquatic vertebrates reflects adaptations to the physical constraints and ecological demands of their environments. For example, streamlined forms, such as those found in sharks, tunas, dolphins, and diving birds, have evolved independently across different vertebrate lineages, highlighting the advantage of this morphology for active swimming in open water; see, e.g., (Kardong 2006; Liem et al. 2001). Similarly, many other body shapes have also arisen convergently, as can be seen from early vertebrates to more derived lineages (Lampart-Kałużniacka and Heese 2000; Li et al. 2020; Nelson et al. 2016; Van Wassenbergh et al. 2015; Wainwright and Turingan 1997). This repeated emergence of the different types of morphologies across diverse taxonomic groups suggests that body shape of vertebrates is evolvable, representing adaptation to distinct environments.

Whereas natural selection in the wild has shaped body forms suited for survival under natural ecological constraints, domestication has introduced a distinct set of selective pressures. A particularly notable case is that of ornamental goldfish, which show a variety of body shapes evolved under aesthetic rather than wild ecological constraints (Le Verger et al. 2024; Ota 2021). Ornamental goldfish were derived from the wild crucian carp (*Carassius auratus*), which show a typical streamlined body shape. Initially, color variations of the single-tail common goldfish were isolated (Fig. 1A-C), and over time, several distinctive morphotypes were selected by breeders and fanciers (Fig. 1D-G) (Chen 1954, 1956; Hervey and Hems 1948; Ota 2021; Ota and Abe 2016; Smartt 2001). These include phenotypes such as popped eyes, bifurcated caudal fins, and the absence of dorsal fins (Chen et al. 2020b; Hervey and Hems 1948; Ota 2021; Ota and Abe 2016; Smartt 2001). Notably, the globular body shape—a rounded and compact form—often emerged concurrently with these mutations in eyes and fins (Asano and Kubo 1972; Kon et al. 2020; Matsui 2006). Thus, this globular morphology is recognized as a defining characteristic of many ornamental goldfish strains (Smartt 2001).

**Fig. 1.**
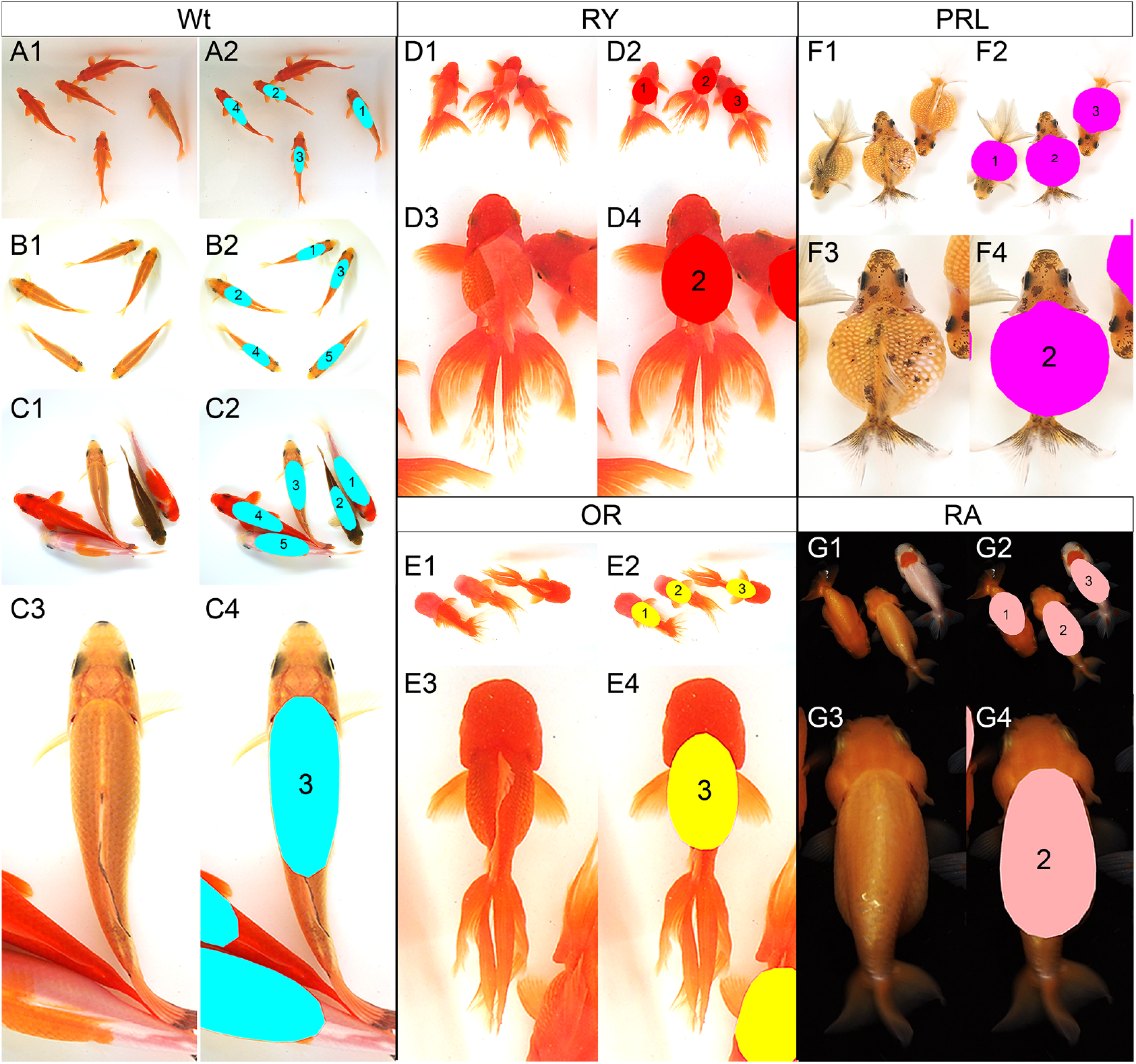
Dorsal view of five ornamental goldfish. A–C. the single-tail common goldfish (wild-type). D. *Ryukin*. E. *Oranda*. F. *Pearl scale*. G. *Ranchu*. The individuals of single-tail, *Ryukin*, and *Oranda* are derived from our previous report (Li et al. 2019; Tsai et al. 2013). The photographs of *Pearl scale* and *Ranchu* are derived from our YouTube channel (https://www.youtube.com/@LAQZL ; see Goldfish EvoDevo series; accessed 2025-06-30). Panel labels ending in odd numbers show the original photographs. Panel labels ending in even numbers show the image with the color-coded region of interest (ROI). The colored ROIs indicate the approximately identified trunk regions. These goldfish individuals are ranging from 5 cm to 15 cm in standard length. The individuals shown in panels A1–B2 were obtained by artificial fertilization and subsequently reared at the Yilan Marine Research Station. The individuals of panels A, B, and C are denoted as Wt-Jp-2019, Wt-Jp-2015, and Wt-Tw-2015, respectively, in Fig. 2 and Table 1. The numbers on each individual correspond to the suffix of the Handle ID in Table 1 (e.g., the “-01” in Wt-Jp-2019-01). Source image available at ark:37281/k5f8z2s55, licensed under CC-BY-NC-SA 4.0, © Kinya G. Ota (2025).

For example, *Ryukin, Oranda, Ranchu*, and *Pearl scale* are ornamental goldfish strains known for their globular body shape (Asano and Kubo 1972; Ota 2021; Smartt 2001) (Fig. 1D–G). Because ornamental goldfish have traditionally been appreciated from a dorsal view (i.e. from ponds and/or opaque fish bowls), their external morphology has been shaped by artificial selection favoring a rounded appearance from a dorsal view. As a result, globular body shapes have become genetically fixed in these populations. Furthermore, if we add an explanation about the nature of this selection, it can be described as follows: this selection appears to have been based primarily on external form, rather than internal phenotypic features. This raised the question whether the globular body shapes seen in different ornamental strains emerged from similar types of modifications, or from distinct internal changes including developmental and tissue reorganization.

**Table 1.** Measured circularity values of five ornamental goldfish.

| Handle ID | Strain | Area* | Perimeter* | Circularity |
| --- | --- | --- | --- | --- |
| Wt-Jp-2019-01 | wild-type | 99357 | 1292.941 | 0.747 |
| Wt-Jp-2019-02 | wild-type | 116019 | 1320.087 | 0.837 |
| Wt-Jp-2019-03 | wild-type | 102129 | 1345.216 | 0.709 |
| Wt-Jp-2019-04 | wild-type | 107958 | 1359.783 | 0.734 |
| Wt-Jp-2015-01 | wild-type | 122932 | 1453.877 | 0.731 |
| Wt-Jp-2015-02 | wild-type | 83594 | 1186.909 | 0.746 |
| Wt-Jp-2015-03 | wild-type | 55259 | 933.59 | 0.797 |
| Wt-Jp-2015-04 | wild-type | 52059 | 948.765 | 0.727 |
| Wt-Jp-2015-05 | wild-type | 71527 | 1136.819 | 0.696 |
| Wt-Tw-2015-01 | wild-type | 210301 | 1988.55 | 0.668 |
| Wt-Tw-2015-02 | wild-type | 145679 | 1662.477 | 0.662 |
| Wt-Tw-2015-03 | wild-type | 185946 | 1790.948 | 0.729 |
| Wt-Tw-2015-04 | wild-type | 204436 | 1950.582 | 0.675 |
| Wt-Tw-2015-05 | wild-type | 227211 | 1981.262 | 0.727 |
| RY-2019-01 | Ryukin | 156483 | 1419.801 | 0.975 |
| RY-2019-02 | Ryukin | 144149 | 1373.393 | 0.96 |
| RY-2019-03 | Ryukin | 156362 | 1439.078 | 0.949 |
| OR-2019-01 | Oranda | 142041 | 1399.848 | 0.911 |
| OR-2019-02 | Oranda | 119059 | 1257.733 | 0.946 |
| OR-2019-03 | Oranda | 122228 | 1303.913 | 0.903 |
| PRL-YT-01 | Pearl scale | 108198 | 1181.011 | 0.975 |
| PRL-YT-02 | Pearl scale | 157832 | 1430.106 | 0.97 |
| PRL-YT-03 | Pearl scale | 123265 | 1261.756 | 0.973 |
| RA-YT-01 | Ranchu | 86237 | 1070.774 | 0.945 |
| RA-YT-02 | Ranchu | 113144 | 1275.322 | 0.874 |
| RA-YT-03 | Ranchu | 83144 | 1095.311 | 0.871 |

\*Area and Perimeter are expressed in pixels as measured directly from digital images.

To address the question posed above, we conducted comparative analyses of representative ornamental goldfish strains (Fig. 1). Recognizing that goldfish fanciers and breeders have traditionally viewed the fish from above and selected for rounder body shapes (Chen 1954; Smartt 2001), we first focused on dorsal morphology and performed shape analysis from this perspective. We then carried out detailed anatomical and histological examinations to identify internal features contributing to the globular body features. We further discuss how artificial selection has historically influenced goldfish morphology and consider whether similar external forms across strains have arisen through shared or distinct developmental mechanisms—potentially representing a case of convergent and parallel evolution at the different phenotypic levels.

## MATERIALS AND METHODS

### Goldfish strains

All goldfish strains used in this study were maintained at the Yilan Marine Research Station (Institute of Cellular and Organismic Biology, Academia Sinica, Taiwan) (Fig. 1). Among them, the wild-type strains were derived from single-tail goldfish originating from Japanese and Taiwanese populations; see also (Ota 2021; Tsai et al. 2013) for a detailed definition of the wild-type goldfish. Progenies used in this study were obtained by artificial fertilization as described previously (Li et al. 2019; Ota 2021; Ota and Abe 2016; Ota et al. 2025). In addition, the *Ryukin, Oranda, Ranchu*, and *Pearl scale* strains were purchased from local ornamental fish distributors in Taiwan. Some of these individuals were directly used for experiments, whereas others were bred to obtain progenies from their parental stocks. For progenies obtained and reared in our laboratory, their origin is specified in the corresponding figure legends.

### Measurement of circularity

Goldfish individuals were photographed from the dorsal side (Fig. 1). The body circularity of goldfish specimens was measured using images captured under controlled conditions. Specimens were recorded against a uniform background (either black or white) using a digital camera or video recording (Olympus; E-M1 Mark III). Images captured from a dorsal view were analyzed to determine the body trunk area and perimeter using the image analysis software Fiji (Schindelin et al. 2012). Details on how the trunk region was defined, as well as the validity of the criteria used, are described in the Results section. These parameters were manually detected. Circularity was calculated based on the detected area and perimeter, using the following formula:

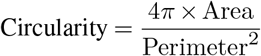

### Anatomical data analysis

After the data acquisition from live fish, the fish were anesthetized with MS-222 (E10521, Sigma). The anesthetized specimens were fixed overnight in 10% formalin. All experimental procedures were conducted in accordance with the animal care and use protocol approved by Academia Sinica (IACUC Protocol 23-10-2072). The fixed specimens were transversely sectioned at the anterior end of the pelvic fin using a microtome blade. The transverse sections were then photographed with a digital camera (Olympus; E-M1 Mark III). From transverse section images, we identified (1) the maximum body width (mbw), defined as the longest distance between the left and right lateral surfaces of the body in the transverse section image, and (2) the internal cavity width (icw), defined as the distance between the left and right boundaries of the body cavity wall. The lateral body wall thickness (lbw)—defined as the combined thickness of the tissue from the skin surface to the body cavity wall on both sides—was calculated by subtracting icw from mbw. In addition, the circularity of the body surface contour and of the coelomic cavity contour was measured from the transverse section images.

### Histological analysis

The 10% formalin fixed specimens from the anatomical data acquisition were dehydrated through ethanol series. After dehydration, specimens were cleared in Lemosol A (5989-27-5, FUJIFILM Wako Pure Chemical Corporation), embedded in paraffin, and sectioned to 5 *µ*m using a microtome (RM2245, Leica). The sections were placed on microscope slides (Platinum-pro, Matsunami). For the conventional histological analyses, the section was deparaffinized with Histo-Clear II (HS-202, National Diagnostics), gradually hydrated with ethanol series, and stained by alcian blue, hematoxylin, and eosin as described previously (Li et al. 2019; Ota et al. 2025). The slide was dehydrated with ethanol series, immersed in Lemosol A, and sealed by coverslip with Entellan new mounting media (107961, Sigma-Aldrich). Histological sections were observed under a microscope, and images were captured focusing on the lateral side of the body wall area. From the histological images, regions containing both muscle and adipose cells were selected, with a focus on areas where adipose cells were densely distributed. Muscle bundles or groups of muscle bundles that were surrounded or adjacent to adipose cells were visually identified and selected. Based on the distribution patterns of muscles and adipose cells, the histological images were scored (see the Results part). The analyses and visualization of the data were conducted by RStudio and SuperPlotsOfData (Goedhart 2021), which is accessible at https://huygens.science.uva.nl/SuperPlotsOfData/.

## RESULTS

To examine the body shape variation among ornamental goldfish strains, we measured the circularity of their dorsal views. Given that the early breeders and fanciers selected their preferred goldfish primarily through visual inspection from dorsal view, we also outlined the body shape of the goldfish and measured the circularity of this outline based on our semi-quantitative visual inspection (Fig. 1). We defined the perimeter of the measured area to include the trunk region, extending from the posterior end of the head to the approximated cloacal area (Fig. 1). Because the cloaca is located on the ventral side and is not directly visible from the dorsal view, its position was approximated using the lateral body width and/or the position of the anal fin. The anal fin, although ventral and normally invisible from above, can be intermittently observed during handling of goldfish, which allowed us to infer the cloaca position. Based on these acquired values, we calculated the circularity of the selected area (Fig. 2 and Table 1). Our approach—though based on visually biased estimations—resembles the way early breeders perceived goldfish and can thus be considered a reasonable proxy for assessing overall body roundness.

**Fig. 2.**
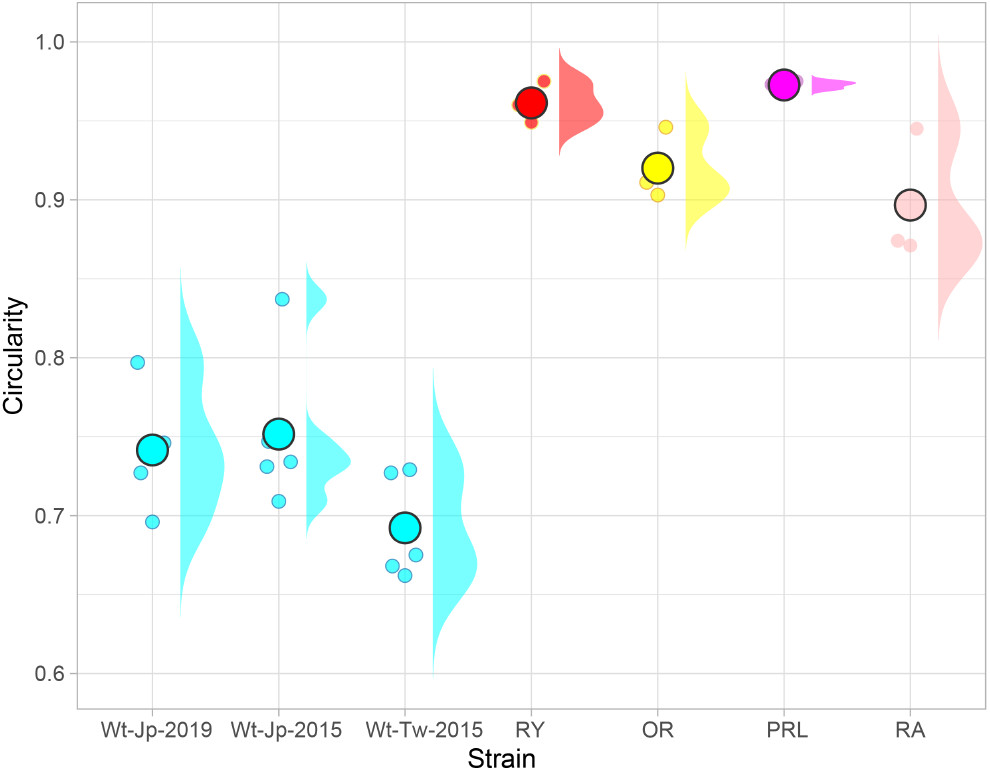
Circularity of five ornamental goldfish. The plot and distribution is based on the image of Fig. 1. The distributions are calculated by SuperPlotsOfData (https://huygens.science.uva.nl/SuperPlotsOfData/). The small dots indicate the circularity of each individual. The large dots indicate the means of circularity. Original data is in Table 1. Source image available at https://n2t.net/ark:/37281/k5f8z2s55, licensed under CC-BY-NC-SA 4.0, © Kinya G. Ota (2025).

From our calculation of the circularity in these strains, it is revealed that the globular body ornamental goldfish, including *Ryukin, Oranda, Pearl scale*, and *Ranchu* strains, showed higher circularities than their wild-type counterparts (Fig. 2). Of those, *Pearl scale* and *Ryukin* goldfish seem to be more circular than the other globular body ornamental goldfish (Fig. 2 and Table 1). We also then compared how the internal anatomical features are different among these ornamental goldfish strains, by observing the section plane of the wild-type, *Ryukin, Oranda, Pearl scale*, and *Ranchu* goldfish strains (Fig. 3A-E). This observation indicated that the lateral body wall thickness is significantly wider in *Pearl scale* goldfish in comparison with the other goldfish strains (the green and blue bars in Fig. 3A-E). *Pearl scale* goldfish also showed a higher ratio of the lateral body wall thickness of body width than the others (Fig. 3F). Crucially, while *Ryukin* and *Pearl scale* individuals showed quite similar circularity (Fig. 2), their anatomical and histological characteristics seem to be highly differentiated from each other. For instance, the transverse section of *Pearl scale* is clearly more circular than that of *Ryukin* (Fig. 3B,D). Moreover, although body size, age, and feeding conditions varied among individuals due to their maintenance in different aquarium tanks, the whitish appearance of the lateral body wall tissue—likely indicating lipid-rich tissue—in the *Pearl scale* goldfish (indicated by black asterisks in Fig. 3D) suggests that this strain possesses distinct morphological and histological characteristics.

**Fig. 3.**
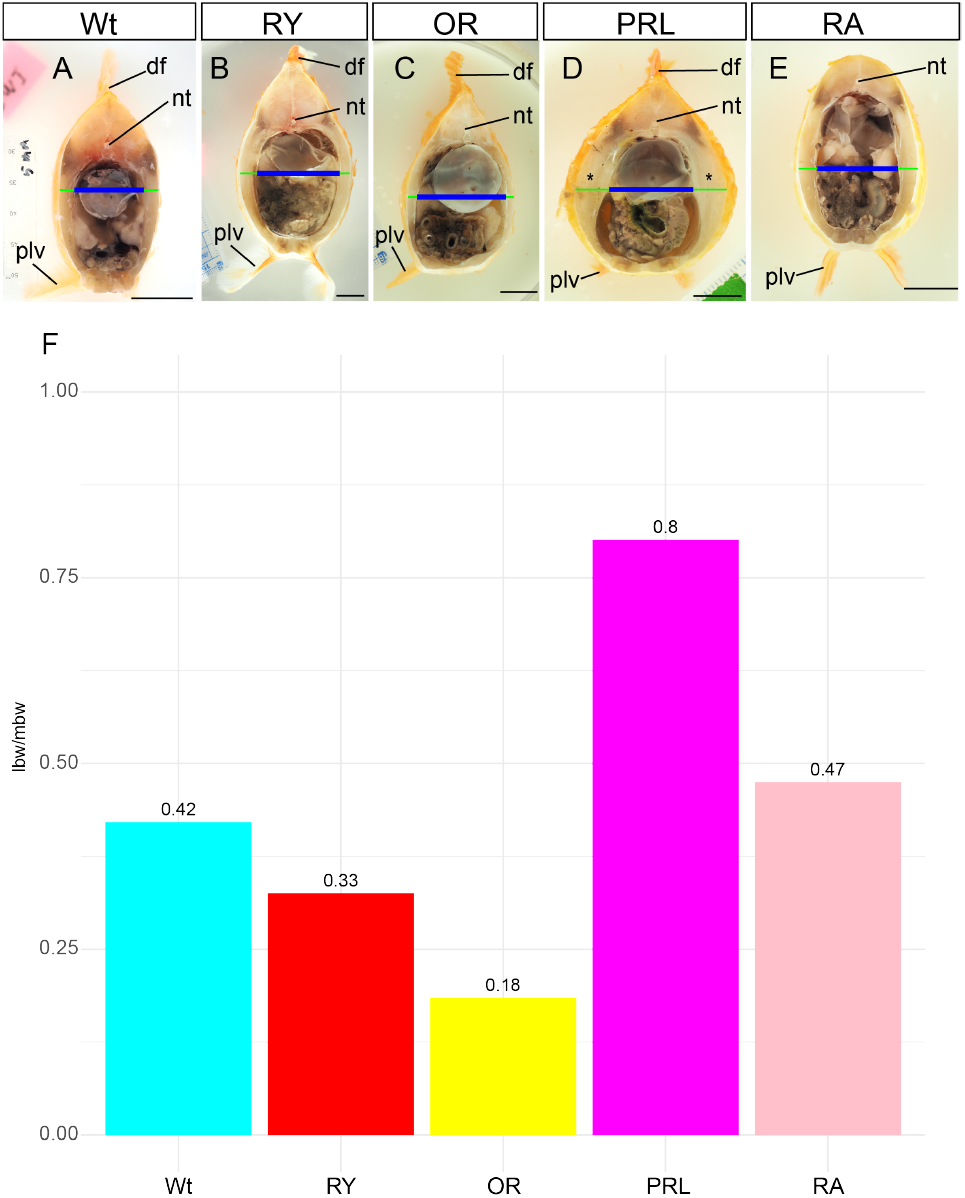
Comparison of the transverse sections in five ornamental goldfish. A-E. Transverse section of the wild-type (Wt), *Ryukin* (RY), *Oranda* (OR), *Pearl scale* (PRL), and *Ranchu* (RA) strain. The blue and green lines indicate the internal cavity width (icw) and the lateral body wall thickness (lbw), respectively. F. Ratio between lbw and maximum body width (mbw). Abbreviations: df, dorsal fin; nt, neural tube; plv, pelvic fin. Scale bars = 1cm. Source image available at https://n2t.net/ark:/37281/k5f8z2s55, licensed under CC-BY-NC-SA 4.0, © Kinya G. Ota (2025).

To further investigate the aforementioned possibility, we obtained a set of data consisting of the circularity of the body and anatomical features of the transverse section plane from seven individuals each from wild-type, *Ryukin*, and *Pearl scale* goldfish (Fig. 4 to 7, and Table 2). The comparison demonstrated a clear trend: both *Ryukin* and *Pearl scale* goldfish have more circular body shapes, and this effect is most evident in the *Pearl scale* strain (Fig. 4). Exceptionally, one of the *Pearl scale* goldfish individuals has a lower circularity than three *Ryukin* individuals (Fig. 4B2, B3, B7, and C7; these panels are indicated by the black and white asterisks). The relatively low circularity observed in this *Pearl scale* individual is likely attributable to its laterally expanded body shape (Fig. 4). These results further confirmed that there are significant differences between the *Pearl scale* and *Ryukin* goldfish strains.

**Table 2.** Measured circularity values of wild-type, *Ryukin*, and *Pearl scale*.

| Handle ID | strain | mbw* | icw* | lbw | lbw / mbw | d-circ** | s-circ** | c-circ** |
| --- | --- | --- | --- | --- | --- | --- | --- | --- |
| 2025-0219-Wt-01 | wild-type | 43.511 | 32.374 | 11.137 | 0.256 | 0.728 | 0.785 | 0.866 |
| 2025-0219-Wt-02 | wild-type | 35.871 | 29.266 | 6.604 | 0.184 | 0.691 | 0.781 | 0.89 |
| 2025-0219-Wt-03 | wild-type | 39.108 | 27.453 | 11.655 | 0.298 | 0.72 | 0.771 | 0.868 |
| 2025-0219-Wt-04 | wild-type | 42.475 | 30.043 | 12.432 | 0.293 | 0.715 | 0.804 | 0.905 |
| 2025-0219-Wt-05 | wild-type | 37.425 | 27.086 | 10.338 | 0.276 | 0.804 | 0.771 | 0.806 |
| 2025-0219-Wt-06 | wild-type | 38.245 | 28.036 | 10.209 | 0.267 | 0.715 | 0.764 | 0.862 |
| 2025-0219-Wt-07 | wild-type | 33.669 | 25.878 | 7.791 | 0.231 | 0.698 | 0.735 | 0.841 |
| 2025-0225-RY01 | *Ryukin* | 47.309 | 33.799 | 13.511 | 0.286 | 0.891 | 0.838 | 0.909 |
| 2025-0225-RY02 | *Ryukin* | 53.029 | 41.698 | 11.331 | 0.214 | 0.961 | 0.887 | 0.97 |
| 2025-0225-RY03 | *Ryukin* | 54.583 | 38.849 | 15.734 | 0.288 | 0.954 | 0.912 | 0.953 |
| 2025-0225-RY04 | *Ryukin* | 40.597 | 32.504 | 8.094 | 0.199 | 0.938 | 0.824 | 0.951 |
| 2025-0225-RY05 | *Ryukin* | 41.957 | 35.029 | 6.928 | 0.165 | 0.891 | 0.853 | 0.959 |
| 2025-0225-RY06 | *Ryukin* | 36.216 | 25.014 | 11.201 | 0.309 | 0.913 | 0.839 | 0.917 |
| 2025-0225-RY07 | *Ryukin* | 38.158 | 28.122 | 10.036 | 0.263 | 0.961 | 0.837 | 0.925 |
| 2025-0225-PRL01 | *Pearl scale* | 105.237 | 64.058 | 41.18 | 0.391 | 0.974 | 0.972 | 0.952 |
| 2025-0225-PRL02 | *Pearl scale* | 82.446 | 54.173 | 28.273 | 0.343 | 0.969 | 0.92 | 0.956 |
| 2025-0225-PRL03 | *Pearl scale* | 84.777 | 45.54 | 39.237 | 0.463 | 0.97 | 0.917 | 0.973 |
| 2025-0225-PRL04 | *Pearl scale* | 74.417 | 47.612 | 26.806 | 0.36 | 0.967 | 0.911 | 0.931 |
| 2025-0225-PRL05 | *Pearl scale* | 71.532 | 49.345 | 22.187 | 0.31 | 0.974 | 0.892 | 0.937 |
| 2025-0225-PRL06 | *Pearl scale* | 66.755 | 49.662 | 17.094 | 0.256 | 0.982 | 0.881 | 0.936 |
| 2025-0225-PRL07 | *Pearl scale* | 64.036 | 40.338 | 23.698 | 0.37 | 0.952 | 0.915 | 0.949 |

\*The values of mbw and icw were obtained from image data and are expressed in pixels. \*\*d-circ, s-circ, and c-circ refer to dorsal-, surface-, and coelom-circularity, respectively.

**Fig. 4.**
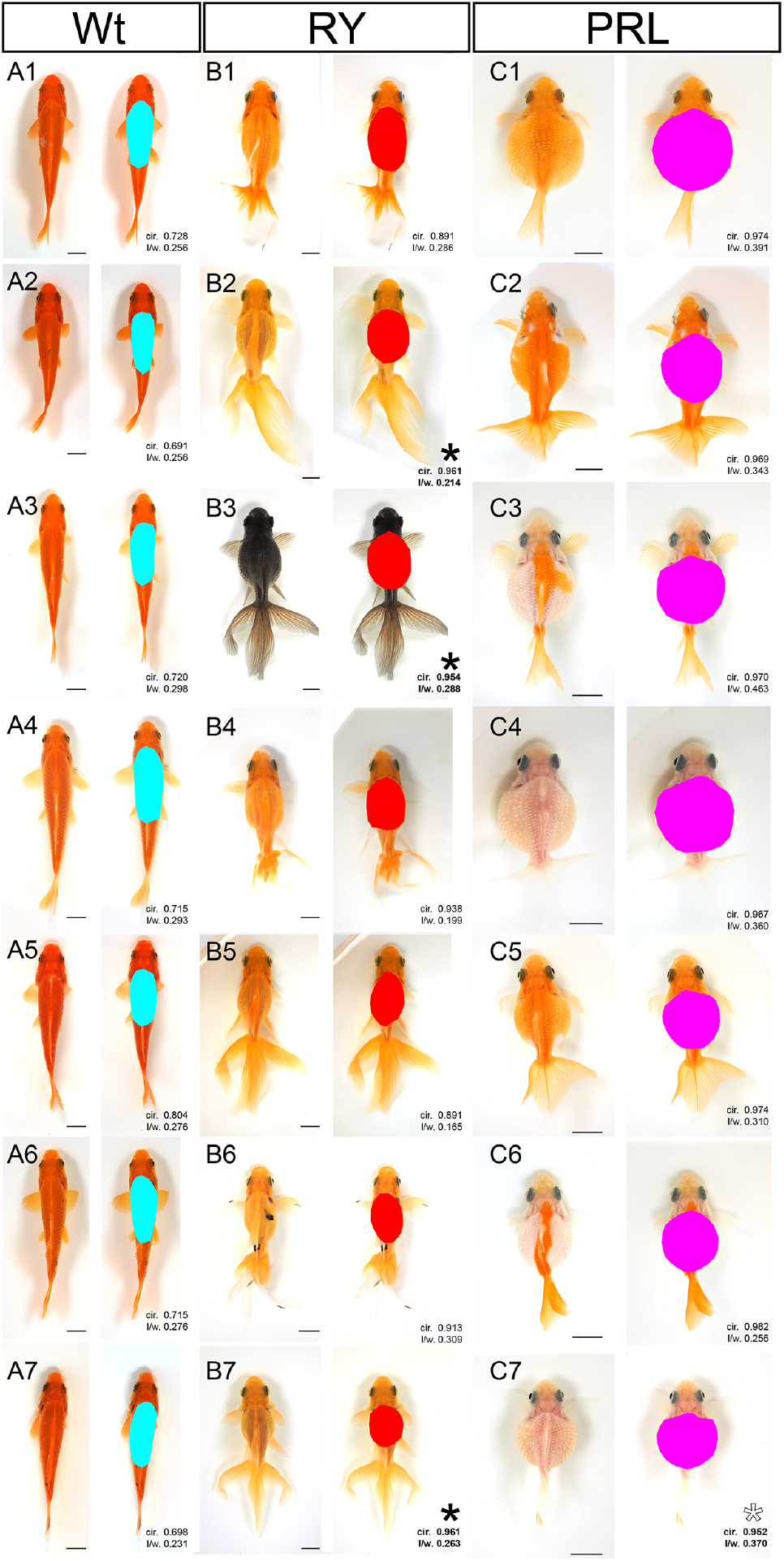
Comparison of circularity between the wild-type, *Ryukin*, and *Pearl scale*. A. Dorsal view of the wild-type (Wt). B. Dorsal view of *Ryukin* (RY). C. Dorsal view of *Pearl scale* (PRL). The light blue, red, and magenta colored areas indicate the approximately identified trunk regions. Black asterisks indicate *Ryukin* individuals showing values approaching those of the *Pearl scale*. White asterisk indicates a *Pearl scale* individual exhibiting values close to those of the *Ryukin*. All individuals shown in this figure were obtained by artificial fertilization and reared in our laboratory. The corresponding data are summarized in Table 2. Scale bars = 1cm. Source image available at https://n2t.net/ark:/37281/k5f8z2s55, licensed under CC-BY-NC-SA 4.0, © Kinya G. Ota (2025).

Next, our results of the transverse section provided consistent and solid results in the differences of the lateral body wall thickness of these examined goldfish individuals (Fig. 5 and 6). The transverse section views in these individuals also indicated that the *Pearl scale* goldfish individuals had a clearly more circular transverse section and showed the largest lateral body wall thickness in comparison with the other strain individuals (Fig. 5). The similar width of the lateral body wall thickness of the wild-type and *Ryukin* strain indicated that although the circularity of the *Ryukin* strain is higher than the wild-type goldfish, this difference stems from the increase of the width of the coelom area in the transverse section view (Fig. 5AB and 6). On the other hand, *Pearl scale* goldfish, although with inter-individual variations, tend to increase the lateral body wall thickness to form the globular body shape (Fig. 6BC).

**Fig. 5.**
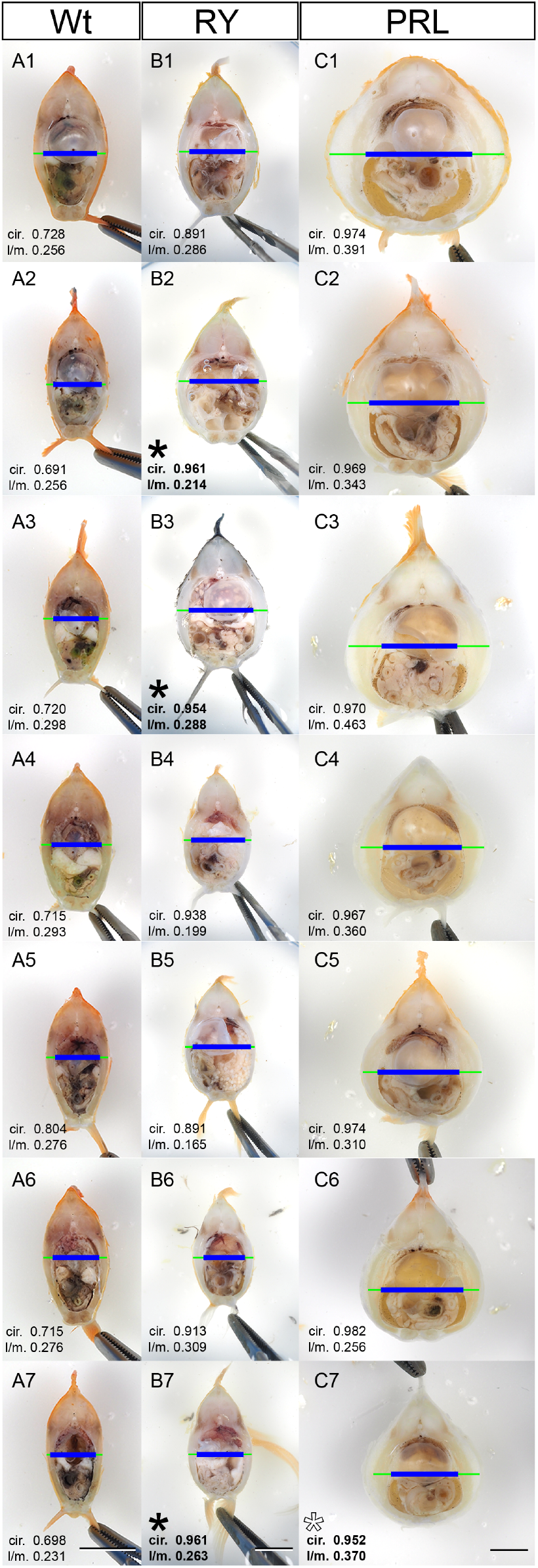
Comparison of the transverse section between the wild-type, *Ryukin*, and *Pearl scale*. A-C. Transverse section of the wild-type (Wt), *Ryukin* (RY), *Pearl scale* (PRL) strains. The blue and green lines indicate the internal cavity width (icw) and the lateral body wall thickness (lbw), respectively. Panel labels correspond to the individual ID in Fig. 4. Numbers of left lower corner in each panel represent circularity and l/m, which is the ratio of the lbw and maximum body width (mbw). White asterisk indicates a *Pearl scale* individual exhibiting values close to those of the *Ryukin*. The corresponding data are summarized in Table 2. Scale bars = 1cm. Source image available at https://n2t.net/ark:/37281/k5f8z2s55, licensed under CC-BY-NC-SA 4.0, © Kinya G. Ota (2025).

**Fig. 6.**
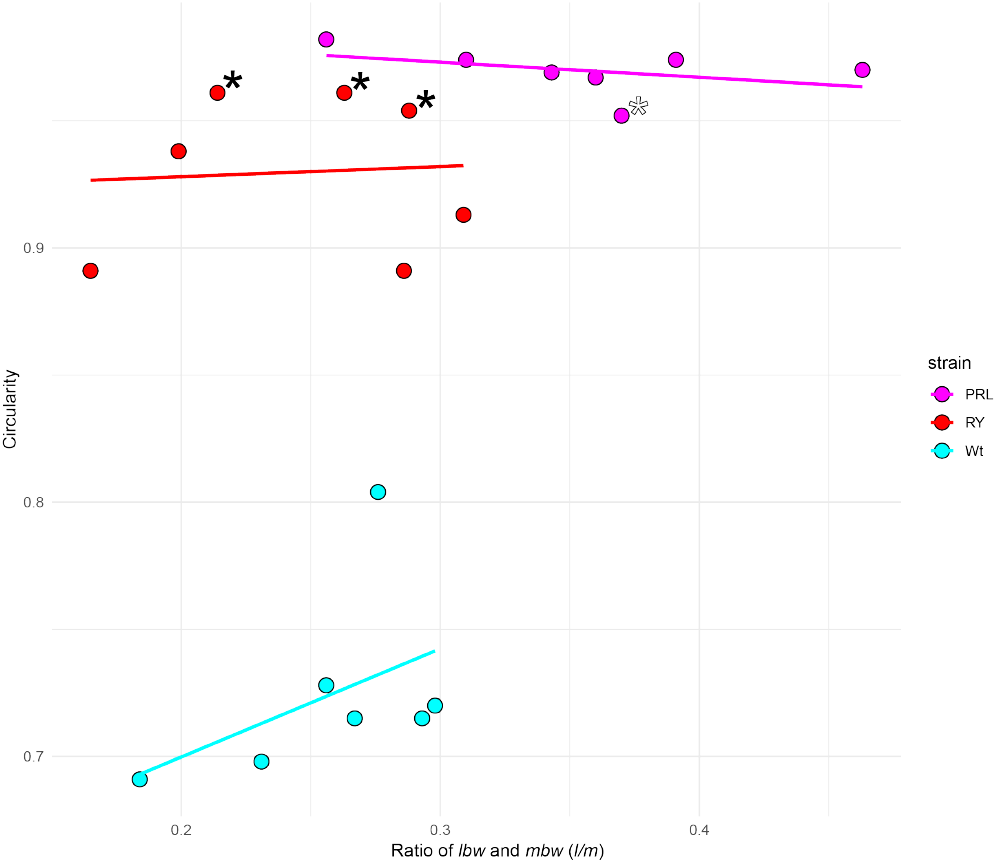
Comparison of circularity and the l/m between the wild-type, *Ryukin*, and *Pearl scale* ornamental goldfish. Light blue, red, and magenta indicate the wild-type, *Ryukin*, and *Pearl scale* goldfish, respectively. The data were derived from Fig. 5 and Table 2.

To clarify further how the body wall and coelom areas changed, the circularities of these structures were measured in transverse section views and compared among the three strains, including wild-type, *Ryukin*, and *Pearl scale* (Fig. 7A). The circularity values of the body surface and coelom in transverse section views indicated significant differences between the wild-type and the two globular body shape goldfish strains. *Ryukin* and *Pearl scale* showed quite similar values, and an increase in the circularity of the coelom occurred in both *Ryukin* and *Pearl scale*, suggesting that the *Pearl scale* goldfish possess a *Ryukin*-like coelom shape while simultaneously exhibiting a thickened lateral body wall (Fig. 6 and 7B). This tendency is also observed in the relationship between body surface circularity in trans-verse sections and dorsal view circularity (Fig. 7C). These results revealed that the globular body goldfish strains are diverged from the wild-type goldfish in the anatomical features. Moreover, while *Ryukin* and *Pearl scale* exhibit comparable anatomical characteristics, their tissue properties appear to be distinct.

**Fig. 7.**
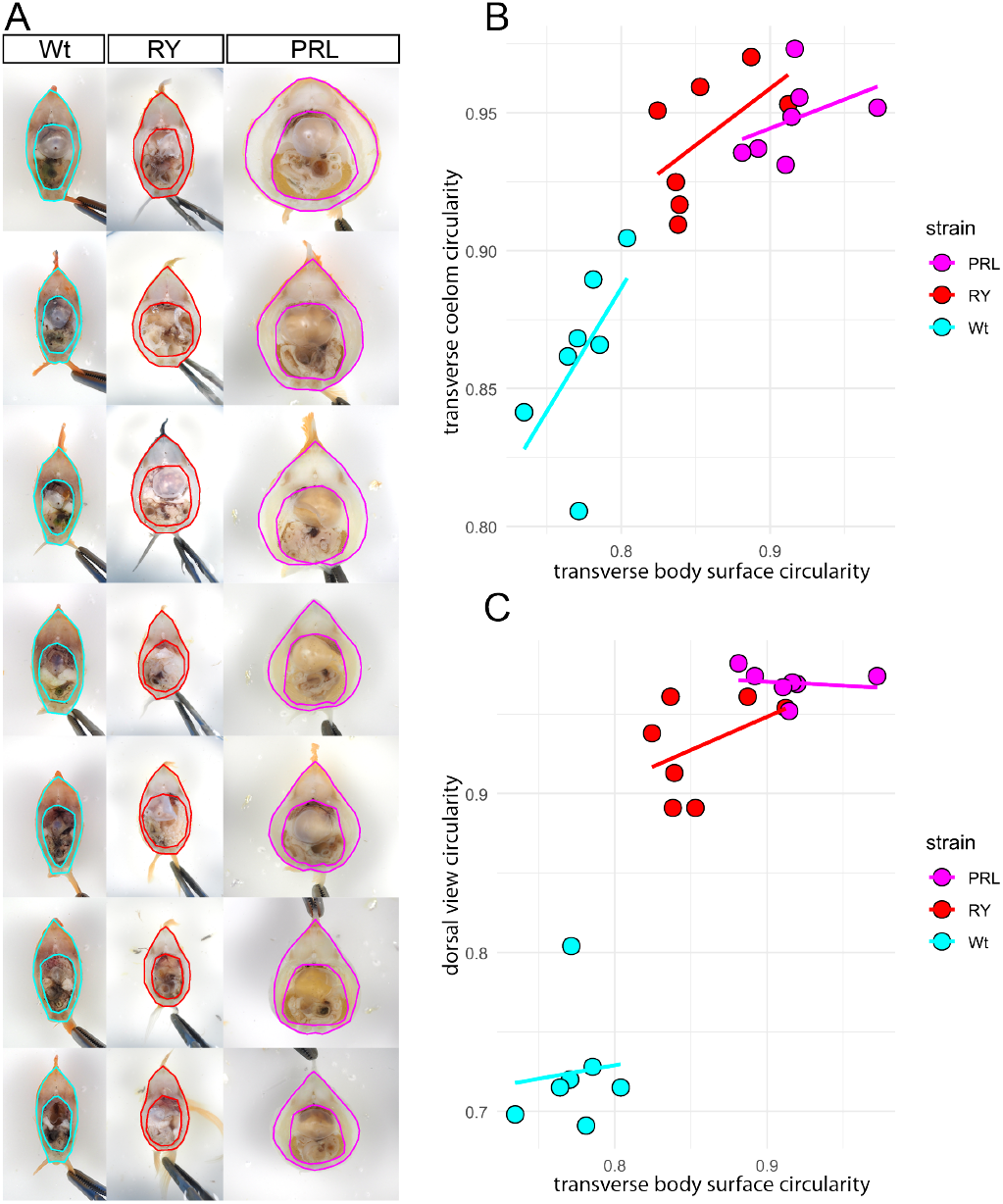
Comparison of circularity of body surface and coelom in transverse section between the wild-type, *Ryukin*, and *Pearl scale* ornamental goldfish. A. Transverse section images from Fig. 5 with outlines of the body surface and coelom traced. B. Plots of body surface circularity and coelom circularity. Light blue, red, and magenta indicate the wild-type, *Ryukin*, and *Pearl scale* goldfish, respectively. The data were derived from Fig. 5 and Table 2 Source image available at https://n2t.net/ark:/37281/k5f8z2s55, licensed under CC-BY-NC-SA 4.0, © Kinya G. Ota

To further examine the similarities and differences at the tissue level between *Ryukin* and *Pearl scale* goldfish, we compared histological features using seven individuals per strain, as described above. We succeeded in observing the lateral area of the trunk region in the transverse section of those individuals at the level of the pelvic fin (Fig. 8). In this observation, we clearly recognized the differences in the ratio between the muscle fibers and adipose tissue in the mid trunk region, as we observed in the transverse hand section views (Fig. 3 and 5).

**Fig. 8.**
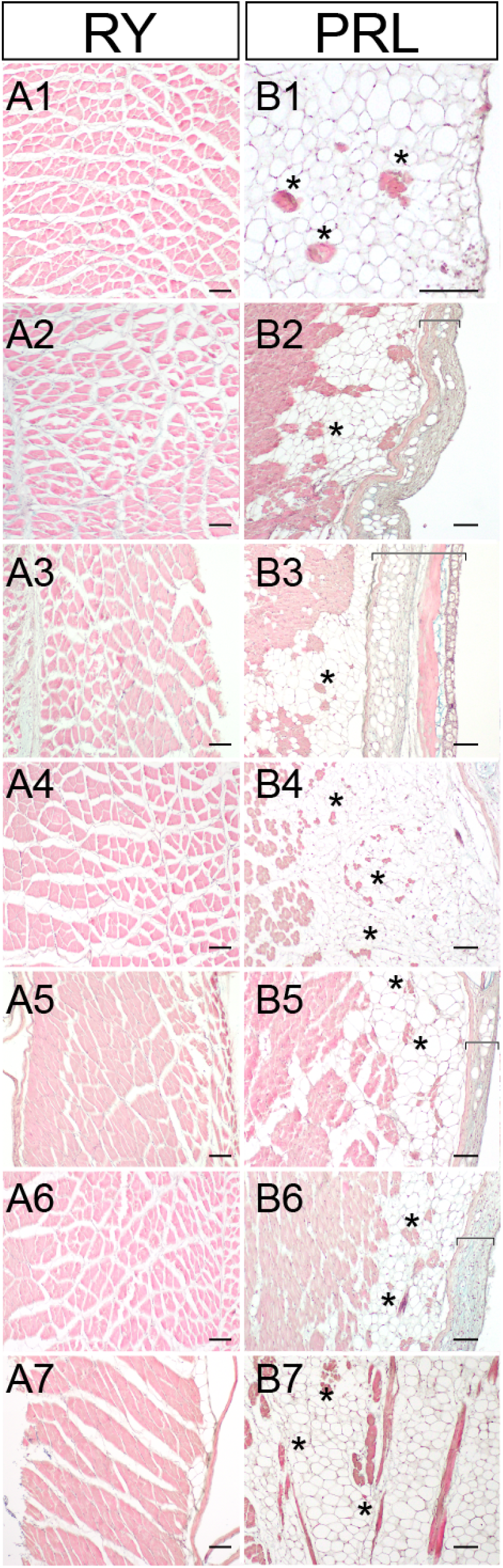
Histological comparison of *Ryukin* and *Pearl scale* goldfish. A–B. Eosin–hematoxylin–Alcian blue-stained sections of the lateral body wall in two globular-bodied ornamental goldfish. Panel numbers correspond to individual IDs shown in Fig. 3. Tissue samples were taken from the region approximately at the level of the pelvic fins, corresponding to the transverse section indicated by the green bars in Fig. 5. Black asterisks indicate muscle fibers surrounded by adipose tissue. The regions containing integumental tissue (epidermis, dermis, scale, and related connective tissues) are labeled in panels B2, B3, B5, and B6 by bracket-shaped bars. Scale bars = 0.1 mm. Source image available at https://n2t.net/ark:/37281/k5f8z2s55, licensed under CC-BY-NC-SA 4.0, © Kinya G. Ota (2025).

The trunk muscle region of the *Ryukin* goldfish showed the conventional muscle tissue (Fig. 8A), i.e. we did not detect adipose tissue surrounding the muscle fibers from our analyzed *Ryukin* individuals (0/7 individuals; Fig. 8A). In contrast, the histological section of the *Pearl scale* goldfish individuals consistently showed muscle fibers surrounded by adipose cells (7/7 individuals; Fig. 8B). Thus, unlike other goldfish that altered anatomical proportions without accompanying histological modifications, *Pearl scale* goldfish appears to have achieved a globular body through both anatomical changes and additional histological modifications involving reorganization of muscle fibers and adipose tissue.

## DISCUSSION

Our comparative analyses revealed that ornamental goldfish can be categorized into three groups in their internal anatomical and histological features: wild-, *Ryukin*-, and *Pearl scale* type (Fig. 9). Although *Ryukin* progeny has a globular body shape, their body wall thickness is quite similar to wild-type, with almost equivalent histological features of the lateral body wall tissue (Fig. 9AB). On the other hand, investigated *Pearl scale* goldfish individuals tend to show a wider lateral body wall (Fig. 6, 8 and 9). We also further revealed that the histological features of *Pearl scale* are highly differentiated from that of the *Ryukin* strain (Fig. 8).

**Fig. 9.**
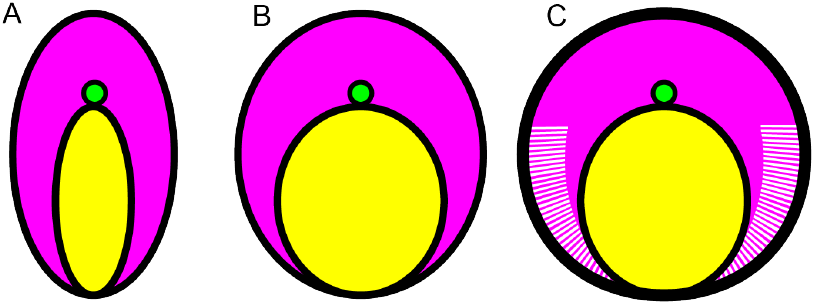
Schematic representation of three different trunk histologies of ornamental goldfish. A. Wild-type. B. *Ryukin*-type, C. *Pearl scale* type. Red, yellow, green, and white areas indicate the relative trunk musculoskeletal area, coelom, neural tube, and adipose tissue. Source image available at https://n2t.net/ark:/37281/k5f8z2s55, licensed under CC-BY-NC-SA 4.0, © Kinya G. Ota (2025).

*Pearl scale*-type ornamental goldfish, along with other related strains, are documented to have been established as early as the Ming dynasty and maintained in ceramic containers traditionally used for fish keeping (Chen 1954). The use of such top-view-oriented containers suggests that morphological selection in ornamental goldfish has historically emphasized dorsal appearance (Fig. 1). From this evidence, it can be inferred that the phenotypic features which can be observed from the lateral side, for example, ratio of the epaxial and hypaxial region, lateral line position, and so on, have not been the initial subject of the selective pressure. Similarly, it is also easily assumed that the internal anatomy, for instance, the volume and circularity of the coelom as well as the lateral body wall thickness, have not been the subject of the attention by the breeders and fanciers (Fig. 3 and 6).

Thus, the currently observed phenotypes are naturally interpreted as a consequence of the selective pressure toward externally visible body shape from the dorsal view, without attention to the internal anatomy and histology. Moreover, it is reasonable to assume that none of the breeders or fanciers have applied any selective pressure specifically to *Pearl scale* goldfish to increase the lateral body wall tissue and its adipose cells. Taken together, *Pearl scale* and other globular body-shaped goldfish might have been under nearly equivalent selective pressures to establish attractive morphological features from the dorsal view. Coincidentally, two distinct histological phenotypic features emerged: an increase of adipose tissue in the lateral body wall of the *Pearl scale* strain, and no such histological modifications in the other globular goldfish like the *Ryukin* strain (Fig. 3 and 6).

To clarify the process underlying the emergence of the globular body shape, we need to further discuss how distinct internal phenotypes could arise, given that all examined goldfish belong to the same interbreeding species, *Carassius auratus*. Throughout breeding history, ornamental goldfish have been extensively interbred by breeders and fanciers with the aim of establishing new strains (Chen 1954; Smartt 2001). The *Ryukin*-type globular body shape, or a tendency toward this morphology, is shared by several modern ornamental goldfish strains, suggesting that interbreeding has occurred among ancestral strains exhibiting the *Ryukin*-type body form. In contrast, the accumulation of adipose tissue appears to be a distinctive morphological feature of the *Pearl scale* (Fig. 3,5 and 9), which warrants further discussion as a unique histological trait.

In fact, strains derived from hybridization with the *Pearl scale* goldfish are well known among breeders and fanciers, and are commonly documented in books and on websites; see Smartt (2001); Yamatokoriyama City Government (2024); Yatomi City Government (2024); Ye and Qu (2017). For example, according to Ye and Qu (2017), several compound variation strains are introduced, such as the *Pearl scale* body type with a cranial hood (as in the *Oranda* strain; see Fig. 1E). However, Ye and Qu (2017) also mentioned that these strains were established only recently, most likely being consolidated as distinct lineages during the 20th century. Taken together, this evidence suggests that, from the perspective of genetic and evolutionary processes, the globular body shape formed through adipose tissue accumulation in the *Pearl scale* may involve a different genetic background from that of the *Ryukin*-type globular body shape.

At present, due to the lack of intensive and extensive genetic or genomic analyses specifically targeting the globular body of ornamental goldfish, we have no convincing data that directly addresses this issue. Nevertheless, the results of genome sequencing analyses of the *Pearl scale* goldfish may provide useful insights (Kon et al. 2020). According to admixture analyses of multiple goldfish genomes, while the genome sequence of the *Ryukin* and *Oranda* goldfish (including its sub-groups) consists of various populations, that of the *Pearl scale* goldfish appears to consist of a single ancestral population (Kon et al. 2020), suggesting isolated genomic features of this goldfish strain. Moreover, the fact that two independent goldfish genome sequencing projects have not investigated any candidate loci for globular body shape suggests that the responsible genetic factors in such a phenotype may not be easily detected (Chen et al. 2020a; Kon et al. 2020). This, in turn, implies that the globular body shape might result from the accumulation of multiple mutated alleles rather than the action of a single major locus. Therefore, the accumulation of multiple variant alleles is considered to have been important in producing the globular body shape, but the manner of this accumulation likely differed between the *Ryukin*-type and the *Pearl scale*-type. The former seems to have arisen through repeated crossbreeding among multiple lineages, whereas the latter appears to have established through a breeding process that emphasized intra-strain crosses to generate a pure line. Thus, it is reasonable to assume that globular body shape is formed in at least two ways in the ornamental goldfish.

To advance this discussion, it is necessary to specify which phenotypic features are clearly dissimilar between *Ryukin* and *Pearl scale*. However, before doing so, we must clarify the methodological limitations of our study. Our morphological comparisons had limitations that precluded accurate quantitative evaluation between *Ryukin* and *Pearl scale*, as the methods relied largely on subjective visual inspection. In addition, the transverse section analyzed was limited to a single level (at the level of the pelvic fin) (Fig. 3 and 5). Given these constraints, it is difficult to rigorously quantify how dorsal-view circularity relates to that of the coelom and body surface in transverse sections of *Ryukin* and *Pearl scale*, even though differences were observed in these values (Fig. 7).

Therefore, further analyses will be necessary to determine, in a purely geometric manner independent of subjective recognition, where the differences in overall globularity lie between *Ryukin* and *Pearl scale*. In other words, breeders, fanciers, and many observers including ourselves and early researchers may perceive that *Pearl scale* appears rounder than *Ryukin* (Smartt 2001), but it remains to be clarified whether this perceived roundness of *Pearl scale* is primarily attributable to the shape of the coelom or to the influence of adipose tissue. A purely geometric assessment of such differences will thus remain a task for future studies.

Beyond the geometric differences of these two strains, it is certain that, at the histological level, *Pearl scale* possesses accumulated adipose tissue not observed in *Ryukin*. Taking this fact into account, for convenience, we describe the evolutionary process as if each strain itself responded to the selective pressures imposed by breeders and fanciers. The *Ryukin* strain specialized in expanding the coelom area and thereby achieved a highly rounded body shape without modifying subcutaneous adipose tissue. By contrast, the *Pearl scale* strain accumulated adipose tissue within the muscle layer in addition to expanding the coelom. Since *Ryukin* and *Pearl scale* exhibit visually similar coelom circularity at the transverse section plane (Fig. 7 and 9BC), *Pearl scale* goldfish may share some anatomical features with non-*Pearl scale* strains. Nevertheless, these two globular body shape types appear to rely on distinct developmental systems for adipose tissue, implying that similar external morphologies can arise through different underlying developmental and histogenesis processes, resembling cases of convergent or parallel evolution in nature.

To avoid unnecessary confusion or debate, we emphasize that our observations are not readily comparable across taxa, unlike streamlined fish morphologies. While such similarities point to a resemblance with convergent or parallel evolution, placing our findings within broader theoretical debates on convergence remains complex. For instance, several voluminous teleost species, including pufferfish, boxfish, and lumpfish, may superficially resemble globular goldfish strains. In reality, some of these voluminous teleosts exhibit modifications in axial skeletal morphology compared with conventional streamlined teleosts, making them superficially comparable to the relationship between globular body shape ornamental goldfish and wild-type forms (Asano and Kubo 1972; Brainerd and Patek 1998; Li et al. 2015). However, their body forms have been shaped by markedly different selective pressures. Thus, although the comparison is not straightforward, our findings invite consideration of alternative approaches that may provide a more appropriate framework for interpreting these cases.

A more fruitful alternative approach, then, is to move beyond purely functional comparisons, and instead examine how morphological convergence can arise alongside divergence at the tissue level. When the focus shifts from viewing morphology in relation to function to considering it in relation to tissue organization, there appear to be cases where more direct relationship can be drawn. For example, our comparison of *Ryukin* and *Pearl scale* goldfish suggests that, within the same species *Carassius auratus*, the same morphological outcome has been produced under similar selective pressures but through additional histological modifications in the *Pearl scale* goldfish lineage. This perspective may allow us to extend the discussion of convergent and parallel evolution. In particular, the sub-cutaneous adipose tissue of the *Pearl scale* goldfish is comparable with several examples.

One striking example is the blubber of marine mammals. Although blubber represents an adaptation to aquatic life—functioning primarily as thermal insulation—and thus differs fundamentally in function from the adipose tissue observed in *Pearl scale* goldfish, the similarity in histological organization is noteworthy. Blubber is typically stratified into three layers: an outer layer containing epidermis, dermis, and adipose cells; a middle layer composed predominantly of adipose tissue; and an inner layer consisting of a mixture of adipose and muscle (Funasaka et al. 2024; Gómez-Campos et al. 2015; Harper et al. 2008; Hashimoto et al. 2015; Montie et al. 2008; Parry 1949). Remarkably, the sub-cutaneous structure observed in *Pearl scale* goldfish closely resembles this three-layered architecture, although the extent of resemblance varies among individuals (Fig. 8B2, B3, B4, B5, B6). This resemblance implies that the adipose tissue layer may possess the greatest evolutionary plasticity among these three layers, and thus may be the most prone to modification under divergent selective pressures. Moreover, similar patterns of adipose–muscle composition are also observed in humans with obesity, in related laboratory animal models, and in meat-producing livestock such as beef cattle and pigs (Claussnitzer et al. 2015; Craig and Moon 2011; David et al. 2016; Den Broeder et al. 2015; Faillaci et al. 2018; Gandarillas and Bas 2009; Green et al. 2018; Landgraf et al. 2017; Li et al. 2023; McMenamin et al. 2013; Park Seung Ju 2018; Pond 1978, 1992; Smemo et al. 2014; Spurlock and Gabler 2008; Tan and Jiang 2024; Visscher et al. 2017; Zang et al. 2018).

Since the conceptual framework regarding the evolution of adipose tissue is still developing (Ottaviani et al. 2011; Pond 1992), further studies are required to enable comprehensive comparative analyses among the examples mentioned above. Despite the functional differences among these cases, remarkably similar histological features are observed. This is largely attributable to the simple and highly conserved layer structures of dermis, adipose tissue, and muscle fibers across vertebrates. Moreover, the recurrent emergence of similar traits in diverse lineages suggests the presence of deeply conserved developmental mechanisms, as also implied by research demonstrating shared molecular pathways of obesity between zebrafish and mammals (Oka et al. 2010). At the same time, difficulties in identifying the genetic basis of subcutaneous adipose accumulation are evident even in established model species (Claussnitzer et al. 2015; Smemo et al. 2014). Such difficulties may partly explain why no specific loci responsible for the globular body shape phenotype in goldfish have yet been reported, despite multiple large-scale genome sequencing projects (Chen et al. 2020b; Kon et al. 2020).

These histologically simple yet genetically complex characteristics may reflect a key aspect of adipose tissue evolution—namely, that traits such as adipose accumulation are highly plastic and shaped by environmental context (Pond 1992). This plasticity has been widely discussed in evolutionary studies of human obesity, where adipose deposition can be either advantageous or disadvantageous depending on ecological and selective conditions (e.g., Qasim et al. 2018; Speakman 2013; Wells 2012). The case of the *Pearl scale* goldfish exemplifies this principle: a globular body shape unlikely to confer fitness advantages in the natural habitat of *Carassius auratus* became highly favored under ornamental selection. Our observations further suggest that this morphology was achieved through at least two developmental modifications—enlargement of the body cavity and extensive sub-cutaneous adipose accumulation—providing an empirical example of how a highly plastic tissue and its developmental trajectory can evolve under artificial selective pressure.

In the present study, our analyses focused only on body-shape–related phenotypes in a limited number of well-established ornamental goldfish strains at the adult stage. Future research should therefore address how globular body formation and adipose tissue accumulation arise during early development. For the *Pearl scale* strain in particular, it will be important to determine whether the massive deposition of adipose cells originates from embryonic modifications or instead emerges later during somatic growth. In addition, developmental coupling between muscle and dermal tissues suggests that the characteristic domed scales of this strain may share a developmental basis with globular body formation, and this possibility warrants further investigation (Hollway et al. 2007; Le Guellec et al. 2004; Smartt 2001; Stellabotte and Devoto 2007). Taken together, these perspectives emphasize the need to link developmental and evolutionary processes that have shaped ornamental phenotypes. We hope that developmental genetic studies of globular-bodied strains, especially the *Pearl scale* goldfish, will deepen our understanding of how selective pressures acting on visually conspicuous morphologies influence histogenesis in highly plastic traits.

## ACKNOWLEDGEMENTS

We are grateful to Wen-Hui Su (SHUEN-SHIN Breeding Farm) and You Syu Huang (formerly of the Aquaculture Breeding Institute, Hualian) for their valuable technical advice on goldfish breeding in Taiwan. We also wish to thank the current and former members of the Yilan Marine Research Station, Institute of Cellular and Organismic Biology: the late Hung-Tsai Lee, Chia-Chun Lee, Chihi-Chiang Lee, and Tsai Han Chuan for their assistance with the maintenance of aquarium systems; Shu-Hua Lee and Sky Wu for occasionally providing brine shrimp when our own supply was lacking; Shi-Chieh Liu for designing aquarium systems; and Chi-Fu Hung, Jhih-Hao Wei, and Fei Chu Chen for administrative support. We are further indebted to Konstantin Khalturin for suggesting comparative approaches with pufferfish. We also wish to thank Qichang Ye for generously donating books relevant to this study. In addition, we thank Kentarou Hirashima and Hiroshi Sugino for valuable feedback on the preprint and for independently confirming several of our observations. We extend our sincere appreciation to Teng Yu-Feng (Big Wind Technology Co., Ltd.), F. L. Lu (YuanLi Instrument Co., Ltd.), and Huang He Nan (YuShing Ltd.) for their assistance in arranging various detailed and often demanding equipment logistics. Finally, we acknowledge the support of the Taiwan Zebrafish Core Facility at Academia Sinica.

## FUNDING

The funding was provided by National Science and Technology Council (Grant No. 112-2311-B-001-033 and 113-2311-B-001 -032 -MY3), Academia Sinica through the Postdoctoral Scholar Program (Grant No. 235g), JSPS KAKENHI (Grant No. JP22K06232), and Takeda Science Foundation (Grant No. 2022036015).

## AUTHOR CONTRIBUTIONS

**Conceptualization**: Kinya G. Ota 太田欽也 (KGO). **Data curation**: KGO, Paul Gerald Layague Sanchez (PGLS). **Formal Analysis**: KGO, PGLS. **Funding Acquisition**: KGO, PGLS. **Investigation**: KGO. **Methodology**: KGO, Chen-Yi Wang 王貞懿 (CYW), Ing-Jia Li 李穎佳 (IJL), PGLS. **Project Administration**: KGO. **Resources**: CYW, IJL, KGO. **Supervision**: KGO. PGLS. **Image acquisition**: KGO. **Writing – Original Draft**: KGO. **Writing – Review & Editing**: KGO, Gembu Abe (GA), PGLS, IJL.

## COMPETING FINANCIAL INTERESTS

The authors declare no competing financial interests.

## AVAILABILITY OF DATA AND MATERIALS

Data underlying the figures and results presented in this manuscript are available in Depositor at https://n2t.net/ark:/37281/k5f8z2s55.

## Notes

https://pid.depositar.io/ark:37281/k5f8z2s55

### Competing Interest Statement

The authors have declared no competing interest.

### Summary of Updates

During a careful re-evaluation of the manuscript, we identified several areas that required clarification or correction. The main updates in this version are as follows: The Abstract was revised to improve clarity and better reflect the main findings. The first paragraph of the Background section was rewritten to provide clearer context for the study. A missing in-text citation for Table 2 was added. Strain abbreviations were added to the panels in Fig. 4, Fig. 5, and Fig. 7 to improve readability. The Discussion section was revised; in particular, the final paragraph was substantially rewritten to clarify our interpretation of the results. Additional contributors were acknowledged, and their names were added to the Acknowledgements section. We believe these updates have improved the clarity and readability of the manuscript. No other changes were made in this version.

